# Altered Tonsillar Microbiome in Children with Down Syndrome and Obstructive Sleep Apnea

**DOI:** 10.64898/2026.05.29.728812

**Authors:** E Woods, D Jones, O Gorden, N Nusbacher, J Kofonow, G Dumont, D N Frank, N R Friedman, B W Herrmann, C Lozupone, M R Verneris

## Abstract

**Background and Objectives:** Children with Down syndrome (DS) have a high prevalence of obstructive sleep apnea (OSA) due to anatomic, neuromuscular, immunological and metabolic factors, yet the contribution of the tonsillar microbiome to airway obstruction in this population remains unexplored. We hypothesized that DS-associated OSA would be associated with a distinct tonsillar microbiome compared to non-DS OSA.

**Methods:** Tonsillar tissue from 22 DS and 18 NDS participants were analyzed by 16S rRNA sequencing. Alpha and beta diversity were assessed using Faith’s phylogenetic diversity and UniFrac distances, respectively, and significantly different taxa were identified with ANCOM-BC and Mann–Whitney testing.

**Results:** Although overall microbial richness and community structure were similar between groups, overweight DS participants demonstrated increased phylogenetic diversity compared to normal-weight DS peers. Taxonomic profiling of the entire patient cohort revealed that in DS tonsils there were selective alterations in key genera with selective depletion of *Haemophilus* and enrichment of *Staphylococcus*, *Rothia*, and *Lactobacillales*. *Haemophilus* abundance correlated positively with tonsil weight in both cohorts.

**Conclusions:** These findings suggest that while global diversity is preserved, specific microbial shifts distinguish the DS tonsillar niche, potentially reflecting altered immune and metabolic environments associated with trisomy 21. Understanding these microbial differences may reveal mechanisms underlying the higher incidence and persistence of OSA in DS and inform targeted therapeutic strategies.

## Introduction

Down Syndrome (DS), caused by trisomy of chromosome 21, is the most common chromosomal disorder worldwide, occurring in approximately 1–2 per 1,000 live births(1). While advances in medical care have extended life expectancy, individuals with DS continue to experience increased morbidity due to a higher prevalence of chronic diseases throughout life, including obesity, hypothyroidism, autoimmunity and sleep-related breathing disorders. Among these, obstructive sleep apnea (OSA) is particularly common, affecting up to 50% of children with DS compared to 1–5% in non-Down syndrome (NDS) populations (2).

OSA is characterized by recurrent partial or complete obstruction of the upper airway during sleep, leading to intermittent hypoxia, fragmented sleep, and systemic inflammation(3). In children, OSA is associated with impaired growth, cardiovascular dysfunction, metabolic disturbances, and neurocognitive deficits(4). In NDS populations, adenotonsillar hypertrophy is a primary cause of airway obstruction, and obesity is a major risk factor for the development of OSA(5).

Increasing evidence has also linked pediatric OSA to dysbiosis of the oral microbiome(6,7). Moreover, tonsils represent a critical immune-microbial interface and altered microbial composition within the tonsillar crypts can drive chronic inflammation and tissue hypertrophy, thereby contributing to upper airway obstruction(8–10). Accordingly, the tonsillar microbiome is significantly dysregulated in OSA patients with lower alpha diversity and distinct microbial composition compared to controls (6,11,12). An overrepresentation of pathogenic genera such as *Haemophilus, Neisseria and Capnocytophaga* and a depletion of *Tannerella, Anaerovorax* and *Halomonas* has also been reported (6,8,11). In the core of hypertrophied tonsillar tissue, *S. aureus*, *H. influenzae*, *N. subflava* and S. *pyogenes* have been reported as being the most prevalent(8).

In DS, OSA is multifactorial, arising from an interplay between anatomic, immunological, metabolic, and neuromuscular factors. Distinct craniofacial features such as midfacial and mandibular hypoplasia, relative macroglossia, and airway narrowing are compounded by generalized hypotonia, leading to increased airway collapsibility(13). This is exacerbated by endocrine abnormalities including obesity, leptin resistance, hypothyroidism, and insulin dysregulation, as well as potential genetic factors including overexpression of chromosome 21 genes (e.g., *DSCR1–4*, *SOD1*) which are linked to metabolic disturbances and oxidative stress, which can amplify inflammation and impair airway muscle tone(14).

Significant differences in the α- and β-diversity of the dental plaque microbiome were found between DS and NDS subjects, which was further influenced by gingivitis. Taxonomic analysis indicated *Corynebacterium*, *Lautropia* were enriched in DS while *Abiotrophia* was depleted when compared to NDS subjects(15). As well, significant differences in the salivary microbiome have been observed between DS and NDS participants, including decreased diversity in DS, characterized by lower abundance of *Alloprevotella*, *Atopobium*, *Candidatus*, *Saccharimonas*, and higher abundance of *Kingella, Staphylococcus, Gemella, Cardiobacterium, Rothia,* and *Actinobacillus*(16).

Because distinct etiologies underlie OSA in DS versus NDS children, we hypothesized that DS-associated OSA would be associated with a distinct tonsillar microbiome compared to non-DS OSA. However, little is known about how trisomy 21 shapes the tonsillar microbial composition in DS. Here we aimed to characterize and compare the tonsillar microbiome in children with and without DS, examining how differences in community diversity and bacterial taxa relate to obesity and OSA risk. Characterizing microbiome alterations may elucidate novel mechanisms underlying airway inflammation and obstruction in DS and inform more targeted therapeutic management strategies.

## Methods

### Ethics Statement

This study was conducted in compliance with the Declaration of Helsinki. Collection of tonsil specimens was approved by the University of Colorado Institutional Review Board (COMIRB 17-2159). As these samples are classified as medical waste, no patient consent was required.

### Sample Collection and DNA extraction

Tonsil specimens were collected on the University of Colorado institutional review board (IRB) number 17-2159. Samples were stored on ice post-surgical extraction and then processed under aseptic conditions in a Class I laminar flow Cabinet. A tissue sample approximately the size of a pea was excised from both tonsils, including the surface and core but avoiding areas of cauterization, and cryopreserved at −20° C prior to DNA extraction.

DNA was extracted using the QIAamp PowerFecal DNA kit (Qiagen Inc, Carlsbad, CA), which employs chemical and mechanical disruption of biomass. Samples were bead-beaten using a MagNA Lyser (Roche Inc, Basel, Switzerland) at 10,000 rpm for 60 seconds

The residual tonsillar tissue was formaldehyde fixed paraffin embedded (FFPE) at the CU Anschutz Gates Institute Histology Core using the VIP5 tissue processor and Histostar embedding unit.

### Bacterial 16S Ribosomal Sequencing

The V4 variable region of the 16S rRNA gene was targeted for sequencing (515F: GTGCCAGCMGCCGCGGTAA, 806R: GGACTACHVGGGTWTCTAAT) and amplified using AccuStart II PCR SuperMix (Quantabio). Construction of primers and amplification procedures follow the Earth Microbiome Project guidelines (www.earthmicrobiome.org). Amplified DNA was quantified in a PicoGreen assay (ThermoFisher Scientfic) and equal quantities of DNA from each sample was pooled. The pooled DNA was sequenced using an Illumina MiSeq v2 2×250 reagent kit on the MiSeq platform at the Anschutz Center for Microbiome Excellence (ACME

### Data processing

The QIME2 microbiome science platform (version 2024.2) was used to process raw paired-end FASTQ files(17). Denoising was then performed with DADA2 to form amplicon sequence variants (ASVs) and a phylogenetic tree was built using sepp. Taxonomy was assigned to ASVs using the RDP Classifier trained on the Silva taxonomic database (release 128) using QIIME 2(18–20). Alpha-diversity was measured using Faith’s PD and beta-diversity was determined by weighted UniFrac distance(21,22).

### Immunohistochemistry

Paraffin blocks were sectioned via microtome at a depth of 5 micron and stained for Hematoxylin and Eosin (H&E), Iron Deposits via Perl’s Prussian Blue and bacteria by Gram Stain at the Gate’s Institute Histology Core. H&E and Prussian blue were imaged at 20x magnification and Gram stain’s were imaged at 60X magnification on an Olympus VS200 Slide Scanner.

### Image analysis

All image analysis was performed using FIJI (https://imagej.net/software/fiji/) version 1.54 q. Body Mass Index (BMI) percentile and BMI Z-Score Calculations:

BMI percentile and BMI Z-score were calculated according to the formulae included in the extended B.M.I. per age growth charts (https://www.cdc.gov/growthcharts/extended-bmi-data-files.htm).

### Statistical analysis

Values were formally compared using Kruskal Wallis or Mann Whitney Rank test. Statistical analysis was performed in QIIME2 or Graph Pad Prism 10.60.

## Results

The demographic characteristics of the cohort are shown in Figure 1. This study included patients with DS (n=22) and NDS (n=18, controls). For the entire cohort, there were 19 males and 16 females (mean age 6.57 years +/− 4.22 years, range 2-18 years). BMI was available for all but 3 (2 DS, 1 NDS) and average BMI was 18.49 +/− 4.15, (range 12.89-30.78). There were no differences in sex or age between patients with DS and NDS (Figure 1A and B). There were also no significant differences between the DS and NDS participants with respect to BMI percentile or BMI z score (Figure 1C-D). As above, the indication for tonsillectomy was OSA and no enrollee had clinical signs of tonsillitis. In prior studies, tonsil size is one of the most important predictors for apnea-hypopnea index (AHI) in preschoolers and school-age children (27). Interestingly, tonsil weight (grams) was significantly smaller in DS compared to NDS participants (Fig. 1E). Tonsil weight was positively correlated with age in DS and NDS (Fig 1F-G) but negatively correlated with BMI only in DS (Fig 1 H-I).

**Figure 1.**
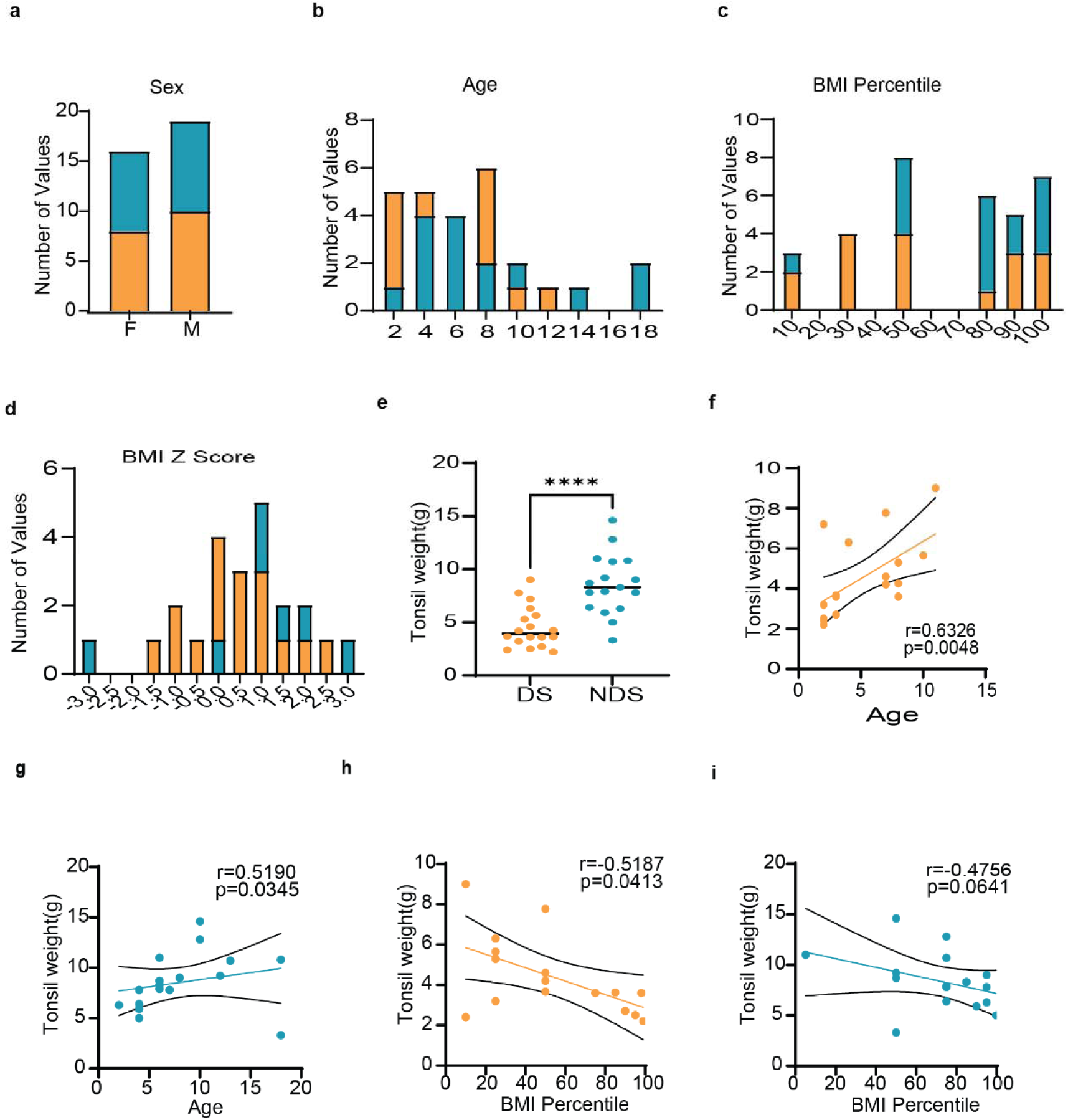
Demographics of Pediatric Obstructive Sleep Apnea Study Enrollees. Frequency Tables showing distribution of **A**. Sex **B.** Age **C**. B.M.I. And **D.** B.M.I. per age percentile of DS (Orange) and NDS (Teal) participants. **E.** Tonsil weight. **F-I** Simple Linear Regression and Spearman’s r showing the relationship between Age and B.M.I. percentile in **F,H** DS **G,I** NDS. Samples were compared by Mann Whitney. **** p <0.0001

Gram staining of tonsil sections in NDS (Fig.2 A-C) revealed that bacterial colonies were primarily localized within tonsillar crypts but also detected in the tonsillar stroma, while in DS bacteria were present within the crypts, the mucosal epithelium and within the tonsillar stroma (Fig.2 D-F). The number of bacterial micro-colonies did not differ between NDS and DS (Fig. 2.G), but in NDS crypt bacterial microcolonies composed of gram-positive bacteria were located adjacent to the crypt mucosal epithelium (Fig.2 B). While in DS, gram-positive bacteria were also visible within the crypt mucosal epithelium and forming microcolonies spanning the crypt epithelium deep into the tonsillar stroma (Fig. 2D).

**Figure 2.**
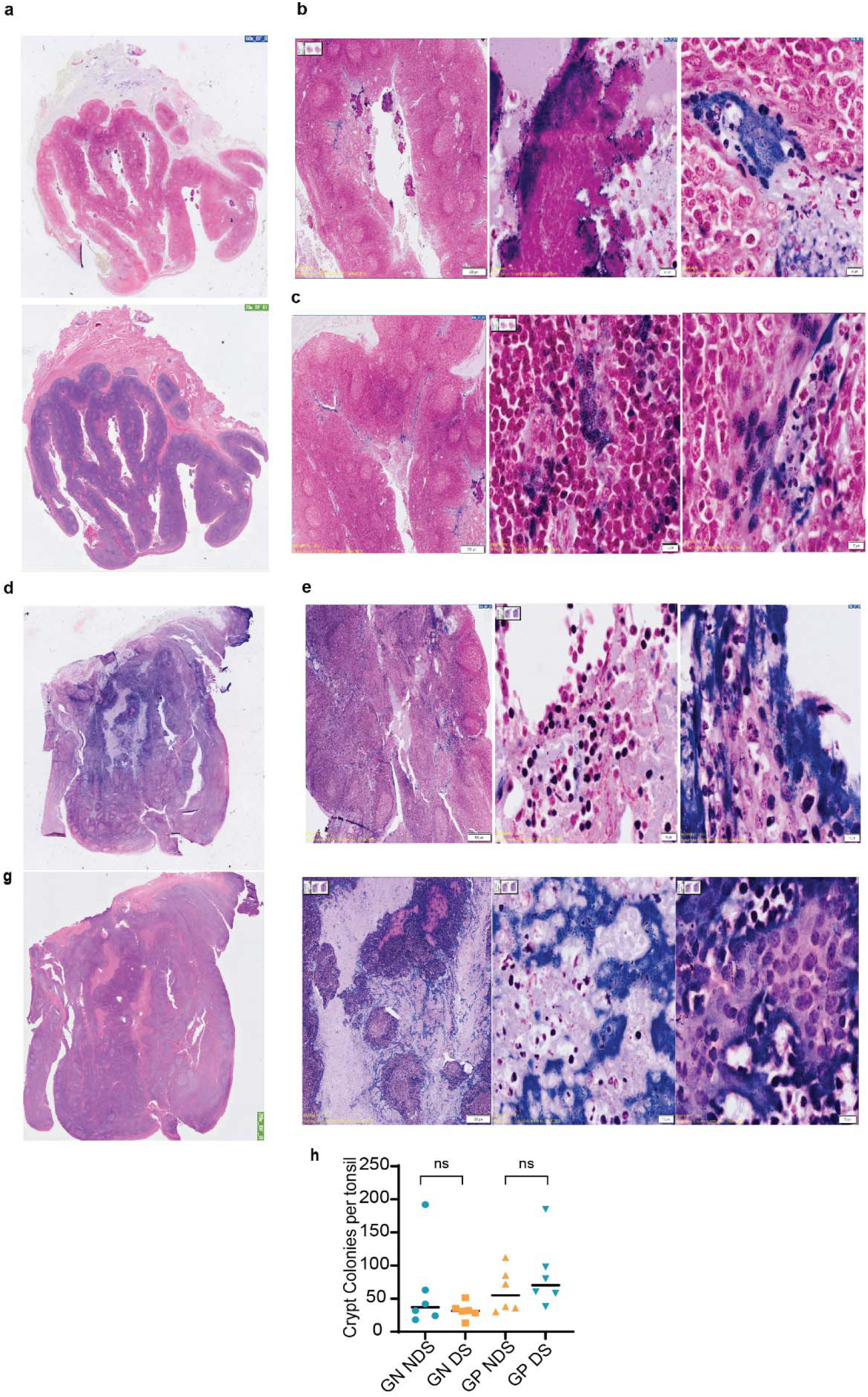
Gram Stain of bacterial microcolonies in NDS and DS Tonsils. **A,D.** Representative tonsil section stained by gram stain (top) or H&E (bottom) **B-C,E-F** overview(left panel) and magnified regions (middle and right panels) of bacterial micro-colonies in the **B,E** Crypt or **C,F** Stroma. **A-C** NDS. **D-F.** DS. Scale Bar: B,E: 500 um (overview) and 10 um (magnified) White arrow: areas of magnification. **G.** Number of Gram negative (GN) or Gram positive (GP) micro-colonies in the tonsillar crypts per tonsil section. Blue-NDS. Orange-DS. H&E: hematoxylin and eosin. GS: Gram stain.

Gram-negative bacteria formed large intra-crypt colonies in NDS that were less abundant in DS (Fig. 2B,D). Collectively, this indicated high level disruption of the gram-negative and gram-positive composition of intra-crypt bacterial niches between NDS and DS, and more translocation of bacteria into the tonsillar stroma in DS OSA tonsils.

### Microbial Diversity of the DS Tonsils

Next, we assessed alpha diversity (i.e. microbial richness and community structure) by measuring Faith’s Phylogenetic Difference (FPD) and evenness respectively within the tonsillar microbiome (21). Analysis of matched tonsils from each participant showed that α- and β-diversity were similar between the left and right tonsils, indicating no significant side-specific differences (Fig S1).

While there were no significant differences in either measure of alpha diversity (FPD and evenness) between DS and NDS children, (Fig. 3.A, B) a positive correlation between BMI percentile and FPD was observed in the DS group, but not in the NDS group (DS - r: 0.5953, p=0.0132; NDS-r:0.2942, p=0.2667, Fig 3C-D). There was no correlation between BMI percentile and evenness for either group (Fig 3E-F). No relationship was present between age or tonsil weight in either group (Fig S2). Interestingly when the dataset was stratified by BMI percentile, a significant increase in FPD (Fig. 3G), but not evenness (Fig. 3H) was observed in the overweight/obese DS subgroup. This suggests that the microbiome structure is impacted when the BMI percentile is above the normal range in DS, but not NDS.

**Figure 3.**
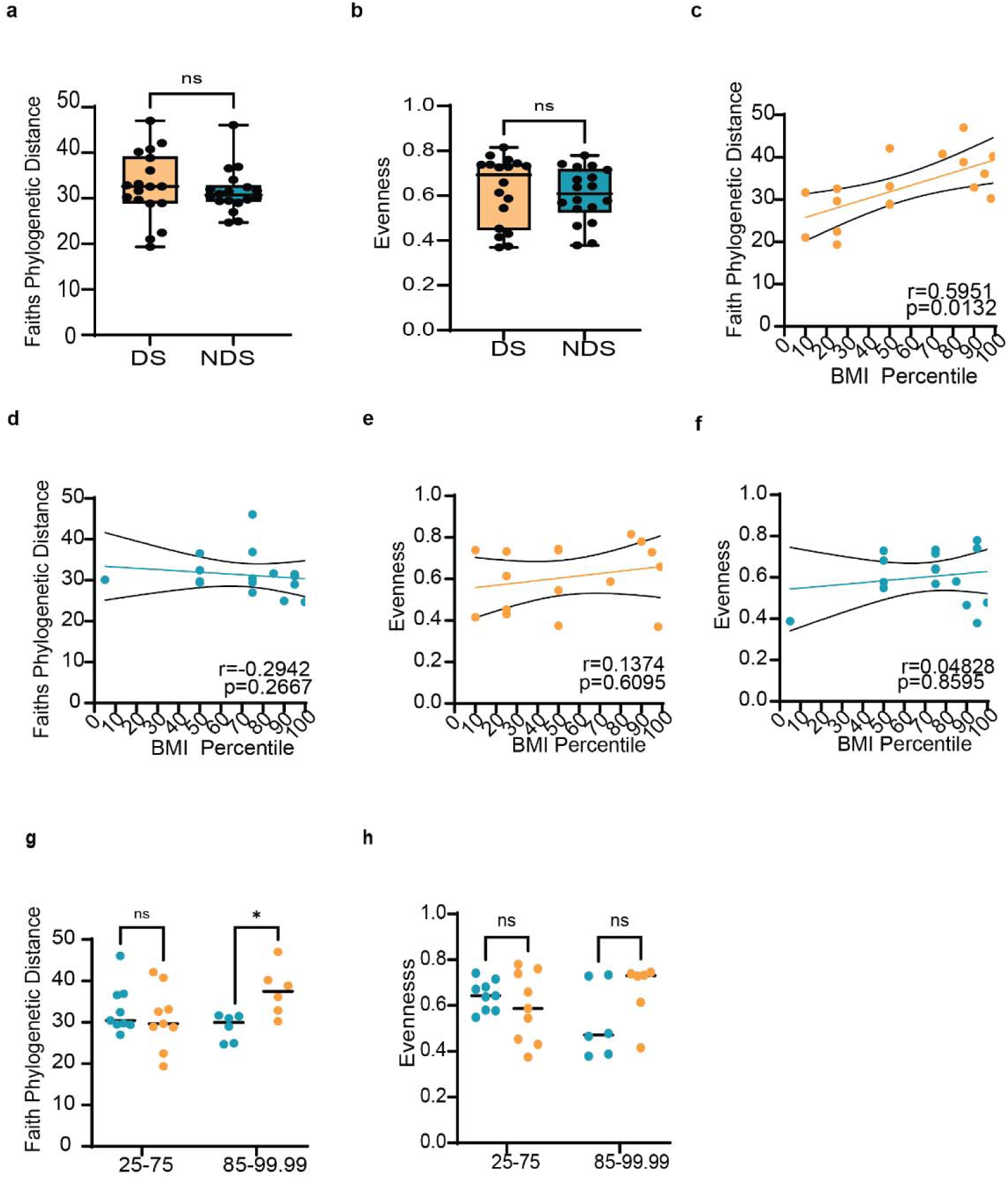
α Diversity of the pediatric OSA tonsillar microbiome is increased in overweight DS but not NDS. **A.** Faiths Phylogenetic Diversity (FPD) **B.** Evenness. **C-F**.Spearman’s r correlation of BMI per age percentile with FPD **(C-D)** and Evenness **(E-F)** in DS (Orange) and NDS (Teal) participants**. G,H.** FPD and evenness in DS and NDS stratified by B.M.I. per age percentile. Samples were compared by Single **(A-B)** or Multiple Mann Whitney Test **(G-H)**, with Holm Sidek correction for Multiple Comparisons at Alpha 0.05. *: p<0.05. ns: Nonsignificant.

To evaluate differences in overall microbial community composition between DS and NDS, beta diversity was assessed by principle coordinate analysis (PCoA) of both weighted and unweighted UniFrac distance (Fig 4)(22). PERMANOVA analysis of both weighted and unweighted UniFrac distances showed no significant differences in overall microbial community composition between DS and NDS groups (Fig 4A-B) or between BMI per age percentile (Fig 4C-D). Collectively, this indicates that the microbial community composition does not significantly differ in OSA between DS and NDS.

**Figure 4:**
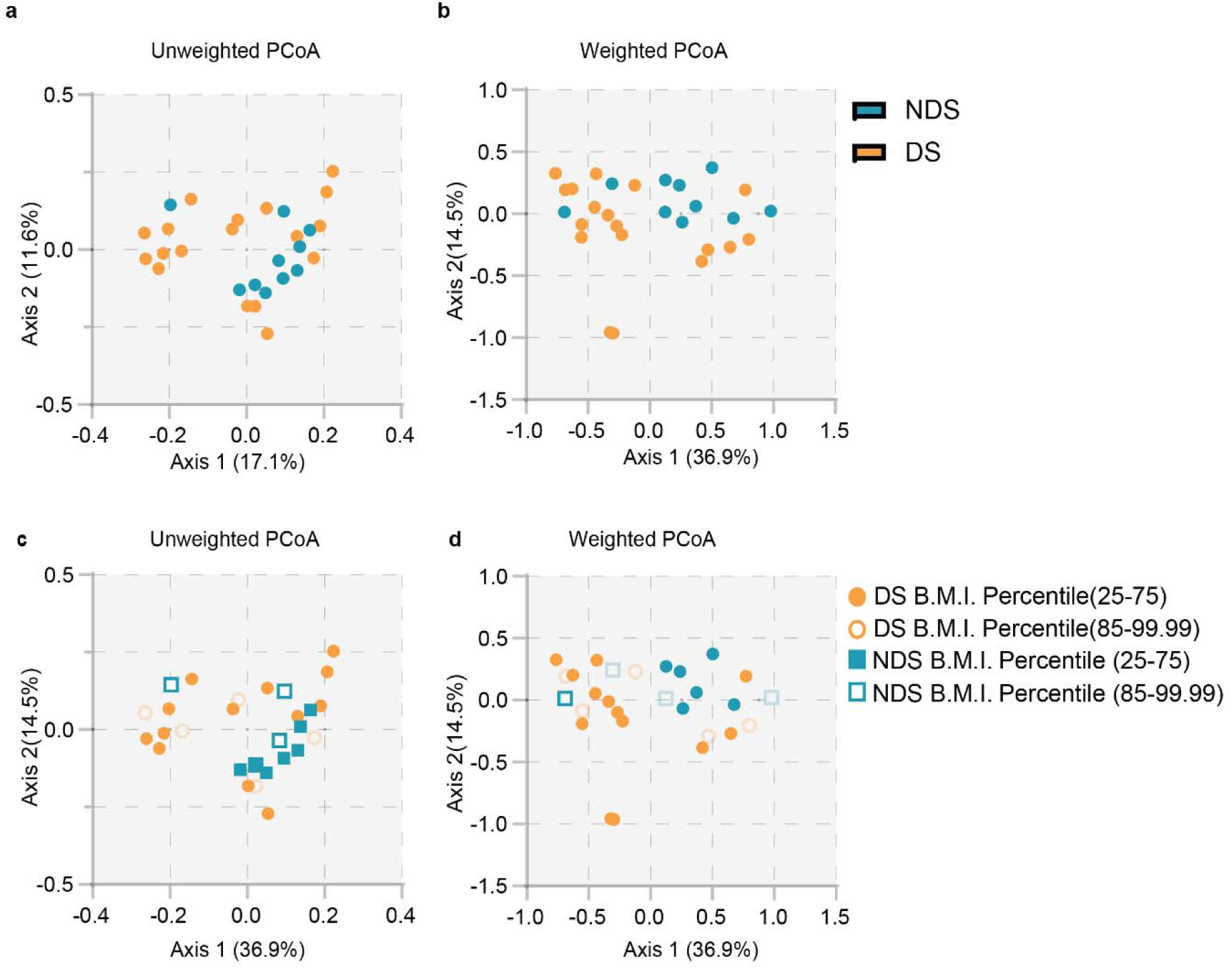
Beta Diversity of the pediatric Tonsillar microbiome in DS and NDS. PCoA analysis of the Unweighted **(A,C)** and Weighted **(B,D)** UniFRAC distance between DS and NDS. Samples were compared via PERMANOVA. Orange: DS, Teal: NDS. Dotted lines indicates origin of X-axis. Axis percentages indicate coverage of Microbiome.

### Taxonomic composition of the DS Tonsillar microbiome

Several bacterial taxa have been implicated in the pathophysiology of pediatric OSA, particularly through their roles in airway inflammation and lymphoid hypertrophy (8,12). We examined the taxonomic composition of the DS tonsillar microbiome at the genus level and compared it to NDS and identified 176 genera across the two groups. Figure 5 shows the individual (Fig 5A) and mean (Fig 5B) taxonomic composition at the genus level of the top 35 most abundant taxa. *Haemophilus* and *Fusobacterium* are common commensals of the upper airway and are enriched in the adenoid and tonsils of pediatric patients with OSA (12). Correspondingly, these were the most abundant taxa in both DS and NDS, followed by *Streptococcus, Leptotrichia, Prevotella, Gemella, Staphylococcus, Parvimonas*, *Neisseria* and *Capnocytophaga*. Differential abundance analysis using ANCOM-BC revealed no significant differences in overall taxonomic composition between DS and NDS (Fig 6).

**Figure 5.**
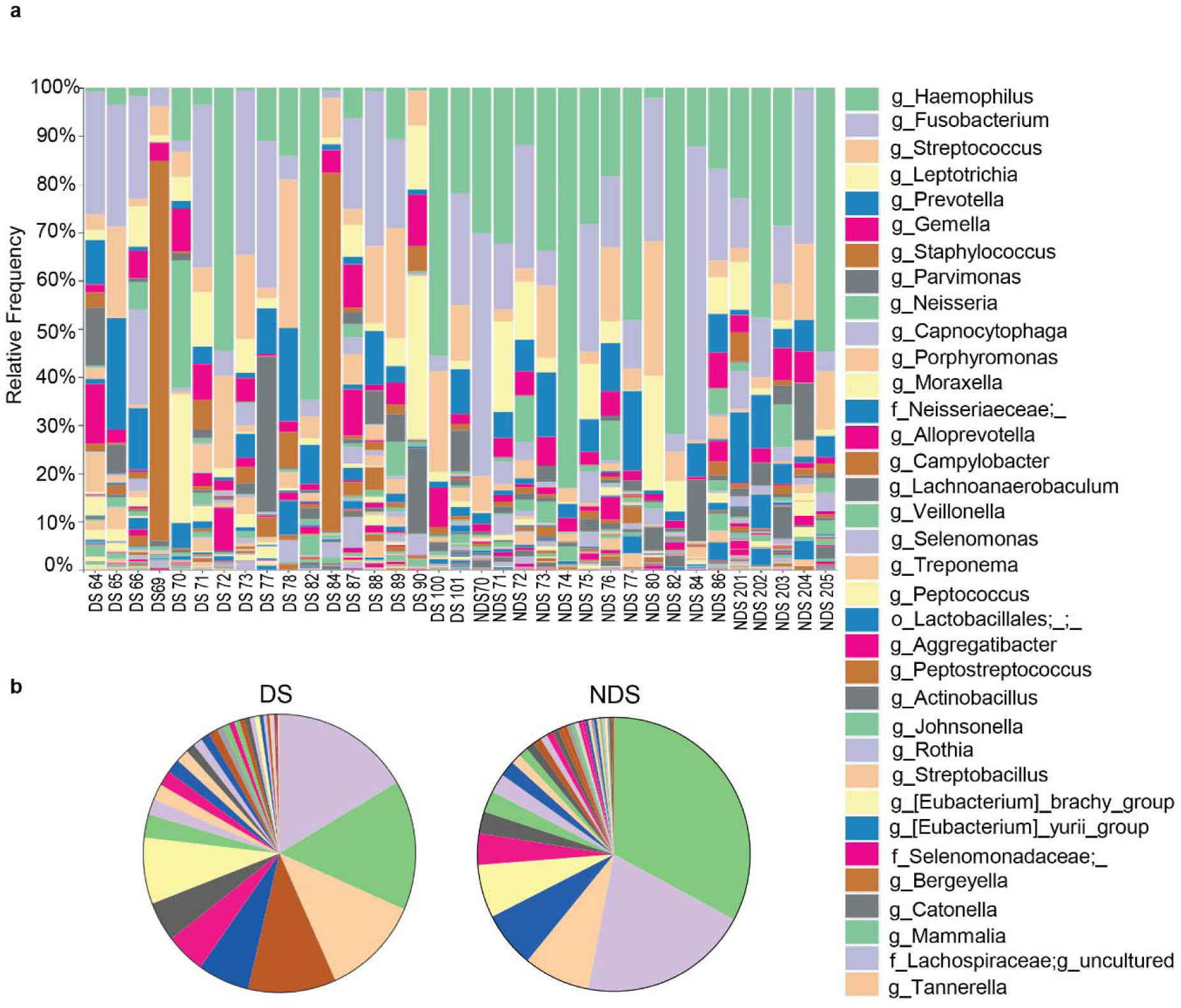
Bacterial community Composition of the tonsils of patients with DS or NDS at the genus level. Relative Abundance at the **(A)** Individual or **(B**) average tonsillar microbiota profiles of enrolled DS and NDS participants. Colors were assigned to the top 35 most abundant organisms detected. (Legend: 35 Most abundant in descending order). g_ : Genus. F_:Familiae. O: Order.

**Figure 6.**
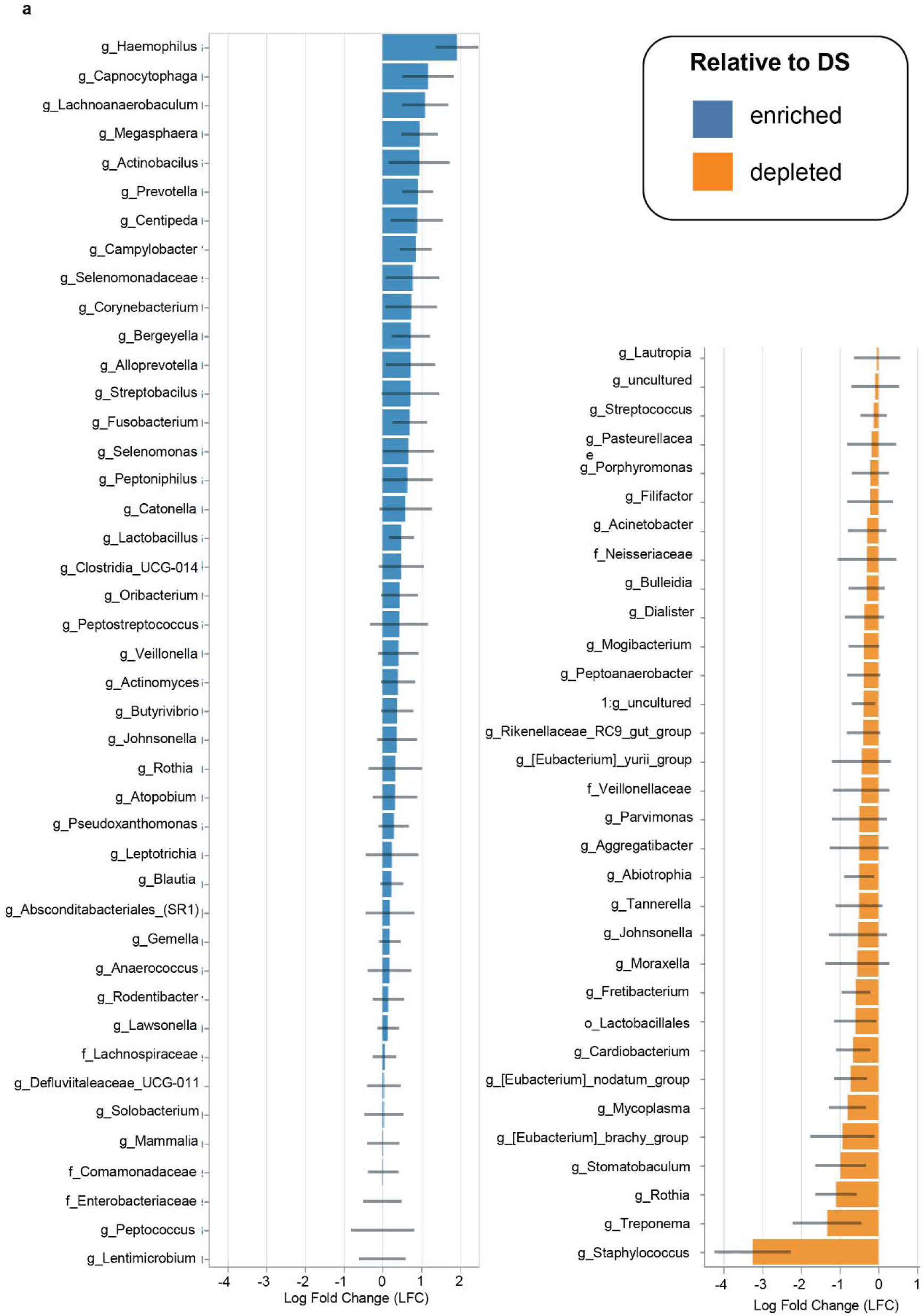
ANCOM-BC analysis of enriched and depleted microorganisms in DS. Data shown is log fold change compared to NDS Reference. Blue: Enriched, Orange: Depleted.

As we hypothesized that DS-associated OSA would be associated with a distinct tonsillar microbiome compared to non-DS OSA, we next performed targeted comparisons on taxa previously associated with NDS OSA. Uusing the Mann–Whitney test, we identified specific taxa that differed significantly between cohorts. *Haemophilus* was significantly decreased in DS (Fig 7A), while *Staphylococcus, Lactobacillales* and *Rothia* were significantly increased compared to the NDS group (Fig 7B-D). As no differences in differentially abundant taxa were found when stratified by BMI per age percentile (Fig. S3), we analyzed the relationship between tonsil weight, age and the differentially abundant microflora in DS and NDS via Spearman’s r correlation analysis. No relationship between *Haemophilus* and age was detected in NDS or between age and any differentially abundant genera. However, tonsil weight was positively correlated with *Haemophilus* abundance in both DS and NDS (Fig 7E-F), which may recapitulate previous findings associating *Haemophilus* with tonsillar hypertrophy (6,8). *Haemophilus* species can rely on metabolic cross-feeding from neighboring taxa. The reduced abundance of *Haemophilus* observed in children with Down syndrome may reflect altered nutrient availability or increased interspecies competition within the tonsillar niche for example, via heme sequestration. Consistent with this, we identified a negative correlation between *Haemophilus* and *Staphylococcus* abundance across pooled NDS and DS but was not significant in individual cohorts, supporting prior evidence that *Staphylococcus* species can inhibit *Haemophilus* (Fig. 7.G).

**Figure 7.**
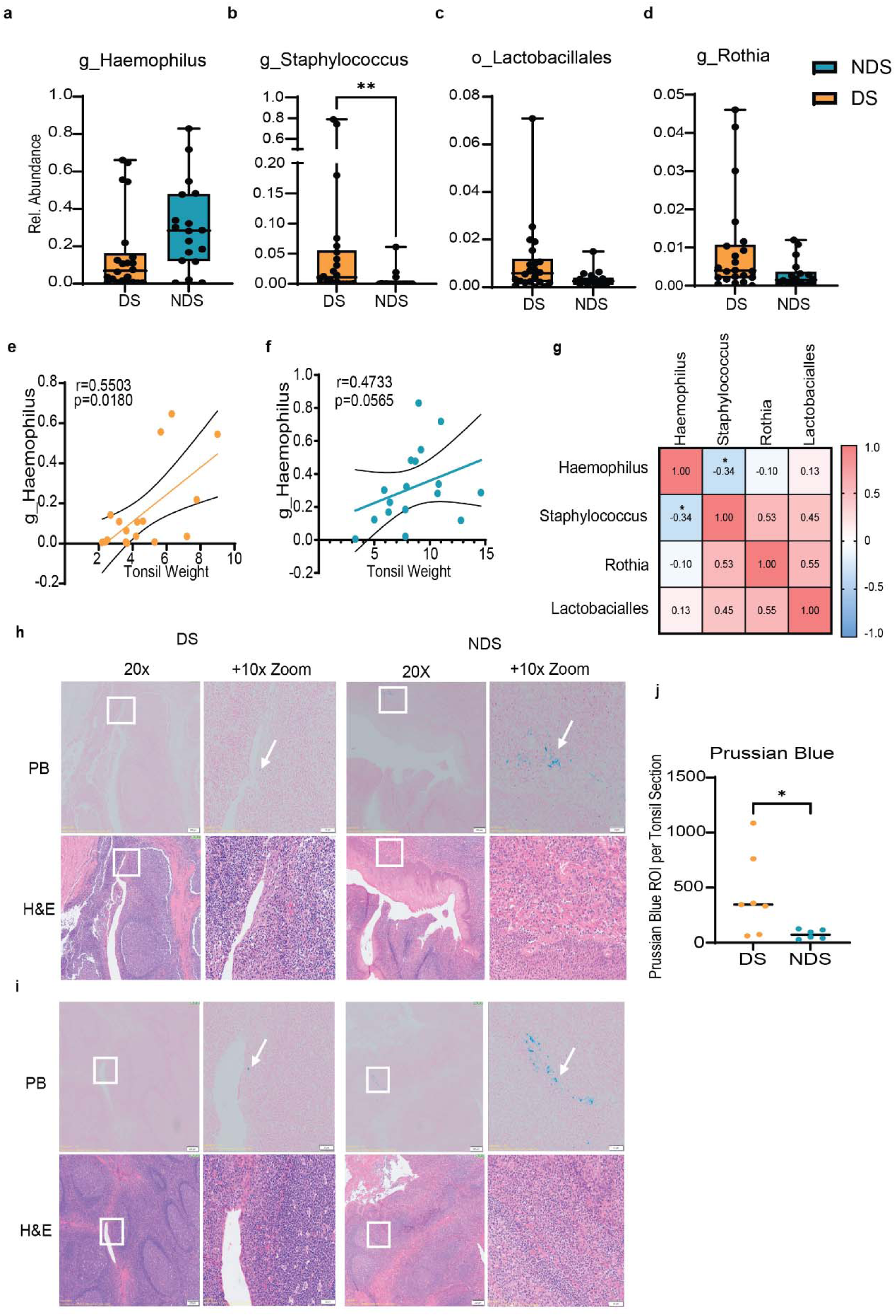
Genus Haemophilus is depleted in the tonsillar microbiome of pediatric DS OSA. Relative abundance of **A**. Genus Haemophilus **B.** Genus Staphylococcus **C.** Order Lactobacialles **D.** Genus. **E.F.** Spearmans Correlation of Haemophilus with Tonsil Weight in DS (Orange) and NDS (Teal)**. G** Correlation matrix of differentially abundant Genera in DS and NDS. Spearman’s r correlation analysis of Cumulative Relative abundance of Haemophilus, Staphylococcus, Rothia and Lactobacialles in all enrollees (DS+NDS). **I-K** Prussian Blue staining of intracellular Iron Deposits in DS and NDS Tonsils. **I,J.** Representative Prussian Blue(PB) and Haematoxylin Eosin (H&E) staining of DS and NDS Tonsils adjacent to **I.** Crypts and **J.** Germinal Centers. 20X Magnification. Left panel: Original, Right Panel: 20X magnification with 10X zoom. Prussian blue: Blue: Iron Deposits, Pink: Tonsillar cells. H&E: Pink: tonsillar cells and parenchyma, Purple/Black: Nuclei. Scale bar original:200um and 10 X zoomed: 50 um. White arrows indicate examples of Prussian Blue regions **K.** Quantification of Prussian Blue Regions in DS Tonsils. Dots indicate total number of Prussian Blue positive regions across one donor tonsil section **A-D,K** Samples were compared by Mann Whitney. *: p<0.05. ns: Nonsignificant. G_: genus. O_: Order

Prussian blue staining suggests increased intracellular ferric iron (Fe³⁺)-deposits in DS tonsils, which may further inhibit *Haemophilus* colonization in DS (Fig 7. I-K). Collectively, these findings suggest the DS tonsillar microbiome in OSA may not have the same taxa enriched as previously identified in NDS OSA, , despite the absence of broad community-level compositional shifts.

## Discussion

Tonsillar dysbiosis has been implicated in adenotonsillar hypertrophy and pediatric OSA in NDS children but has not been studied in DS. Because of the high prevalence of OSA and immune dysregulation in DS, we hypothesized that children with DS exhibit distinct tonsillar microbiome profiles compared with NDS children, with these differences potentially playing a foundational role. This study represents the first characterization of the tonsillar microbiome in DS children undergoing tonsillectomy for OSA compared with NDS.

Despite similar BMI, the age of the DS group trended lower then the NDS group. Correspondingly, Tonsil weight was positively correlated with age across the cohorts, and DS tonsils were significantly smaller than NDS participants.Nevertheless, while adenotonsillar enlargement is a major etiological factor in the pathophysiology of OSA in NDS children , additional variables aside from tonsillar hypertrophy may also contribute to airway obstruction in DS. Indeed in DS, OSA can persist even after adenotonsillectomy (28). Craniofacial and upper airway abnormalities, including midface hypoplasia, macroglossia, glossoptosis, and generalized hypotonia are well defined in DS (29). Co-morbidities for OSA, including obesity, hypothyroidism, and gastroesophageal reflux are also more prevalent in DS and can exacerbate upper airway narrowing and inflammation, amplifying the risk of OSA in DS even in the absence of hypertrophic lymphoid tissue. However, an important limitation of this study is that we did not evaluate non OSA DS and NDS tonsils.

While microbiome profiling revealed no overall differences in alpha diversity (FPD and evenness) or beta diversity (weighted and unweighted UniFrac) between DS and NDS groups, indicating a similar microbial richness, diversity and community structure between DS and NDS, we did observe that FPD was significantly elevated in DS participants when stratified into normal or overweight groups using BMI per age percentile (normal: 15-75 percentile and overweight: >85 percentile). This is intriguing, as obesity is a well-established OSA risk factor and has been linked to striking dysbiosis of the gut microbiome(30–32). Moreover, several studies in NDS people have demonstrated significant differences in oral microbiome composition between normal-weight individuals and those with obesity, which paralleled changes in the obesity associated microbiota of the gut, highlighting the physiological link between the oral and gut microbiome (30–32). While the gut microbiome is dissimilar in NDS and DS and has been correlated to cognitive impairment in DS, the impact of obesity has not been defined in DS (33). Whether changes in the tonsillar microbiome also reflect changes in the gut microbiome in DS and the relationship to cognitive disability and obesity related co - morbidities is an important avenue of future research. However, the necessity of grouping overweight and obese subjects due to low sub-group recruitment means we cannot yet define a “dose-response” relationship between BMI and FPD in DS. Future studies with stratified recruitment are essential to determine if the microbial impact of BMI is linear or if a specific threshold triggers the observed changes in diversity.

Gram stains revealed considerable translocation of gram-positive and gram-negative bacteria from the crypt mucosal epithelium and into the tonsillar stroma. Gram negative micro-colonies were also less frequent in DS crypts compared to NDS. At the genus-level taxonomic analysis identified significant compositional shifts, with *Haemophilus* markedly depleted and *Staphylococcus* and *Rothia* enriched in DS tonsils. Indeed *Haemophilus* was detectable in DS ,which is consistent with prior studies in DS oral flora, it was significantly less abundant compared to NDS OSA, where it was the most abundant taxa identified (16). The smaller tonsil size observed in DS, alongside decreased *Haemophilus* abundance, contrasts with prior reports in NDS OSA where *Haemophilus influenzae* was linked to tonsillar hypertrophy and chronic inflammation (34–36) perhaps underscoring the additional contribution of upper airway abnormalities to OSA in DS. A notable study limitation was the discrepancy between our targeted Mann-Whitney analyses and the ANCOM-BC. While Mann-Whitney identified significant differences in specific organisms previously implicated in pediatric OSA, this warrants further investigation in larger cohorts as our limited sample size likely limited statistical power for ANCOM-BC.

*Haemophilus has a* species level requirement for exogenous heme (X factor) and nicotinamide adenine dinucleotide (V factor) and hence are dependent on metabolic cross-feeding from neighboring taxa (37–39). In DS, reduced abundance of *Haemophilus* may reflect altered nutrient dynamics or interspecies competition within the tonsillar niche (39–42). In this context, we observed a negative correlation between *Haemophilus* and *Staphylococcus* abundance, which supports prior evidence that *Staphylococcus sp.* can inhibit *Haemophilus* growth through heme sequestration and production of antimicrobial metabolites (43,44). The enrichment of *Staphylococcus* in DS could suggest competitive displacement potentially exacerbated by altered immune or metabolic conditions associated with trisomy 21 (45).

Notably, emerging data implicate chronic interferon (IFN) signaling and myeloid dysfunction (including neutrophil and monocyte chemotaxis, phagocytosis, and oxidative burst) in DS (46–49). These likely contribute to impaired bacterial clearance and iron sequestration within macrophages (50–52). In support of this, Prussian blue staining shows increased ferric iron (Fe³⁺)-laden deposits in DS tonsils, potentially limiting extracellular heme availability and further disadvantaging *Haemophilus* growth (41). These findings highlight a potential interplay between host iron metabolism, immune regulation, and microbial ecology in DS.

## Conclusion

Collectively, our results indicate that while the overall microbial diversity of the tonsillar microbiome is preserved in DS, its composition is significantly altered, favoring genera associated with inflammation and reduced colonization by *Haemophilus*. These ecological shifts may reflect systemic features of trisomy 21 and could contribute to the unique pathophysiology of OSA in this population. Further investigation using species-level resolution will be essential to determine causality between microbiome dysbiosis and identify interventions for OSA in DS.

## Acknowledgements

We would like to thank all self-advocates with Down Syndrome and their families for taking part in this study. We’d also like to thank the personnel from the Anschutz Center for Microbiome Excellence (ACME), the University of Colorado Anschutz Medical Campus Advanced Light Microscopy Core and the Gate’s Institute Histology Core. Imaging was performed in the Advanced Light Microscopy Core facility of the NeuroTechnology Center at the University of Colorado Anschutz Medical Campus, which is supported in part by Rocky Mountain Neurological Disorders Core Grant (P30NS048154) and by Diabetes Research Center Grant (P30 DK116073). This work was funded under R01HL155691-01.

## Author Contributions

**EW** and **DJ** collected data, carried out initial analyses conceptualized and designed the study, drafted the initial manuscript, and critically reviewed and revised the manuscript. **OG** collected data, carried out the initial analyses and critically reviewed and revised the manuscript, **DF,NN**, **GD** and **JK** collected data, carried out the initial analyses. **BH** and **NF** collected Tonsillectomy Samples and critically reviewed and revised the manuscript. **MRV** and **CL** conceptualized and designed the study, coordinated and supervised data collection, and critically reviewed and revised the manuscript for important intellectual content. All authors approved the final manuscript as submitted and agree to be accountable for all aspects of the work.

## Data availability statement

The datasets generated and analyzed during the current study are available in the NCBI Sequence Read Archive (SRA) repository under BioProject ID PRJNA1441615 (https://dataview.ncbi.nlm.nih.gov/object/PRJNA1441615). Additionally, datasets will be made accessible via the INCLUDE Data Hub.”

## Additional Information

### Conflict of Interest Disclosures (includes financial disclosures)

The authors have no conflicts of interest to disclose.

### Role of Funder/Sponsor (if any)

The NIH had no role in the design or conduct of this study.

### Abbreviations

ACME: Anschutz Center for Microbiome Excellence
ASVs: amplicon sequence variants
BMI: Body Mass Index
DS: Down Syndrome
FPD: Faith’s Phylogenetic Difference
FFPE: formaldehyde fixed paraffin embedded
H&E: Hematoxylin and Eosin
IFN: Interferon
IRB: Institutional Review Board
NDS: Non Down Syndrome
OSA: Obstructive Sleep Apnoea
PCoA: principle coordinate analysis

