## Supplemental figures for "Altered Tonsillar Microbiome in Children with Down Syndrome and Obstructive Sleep Apnea"

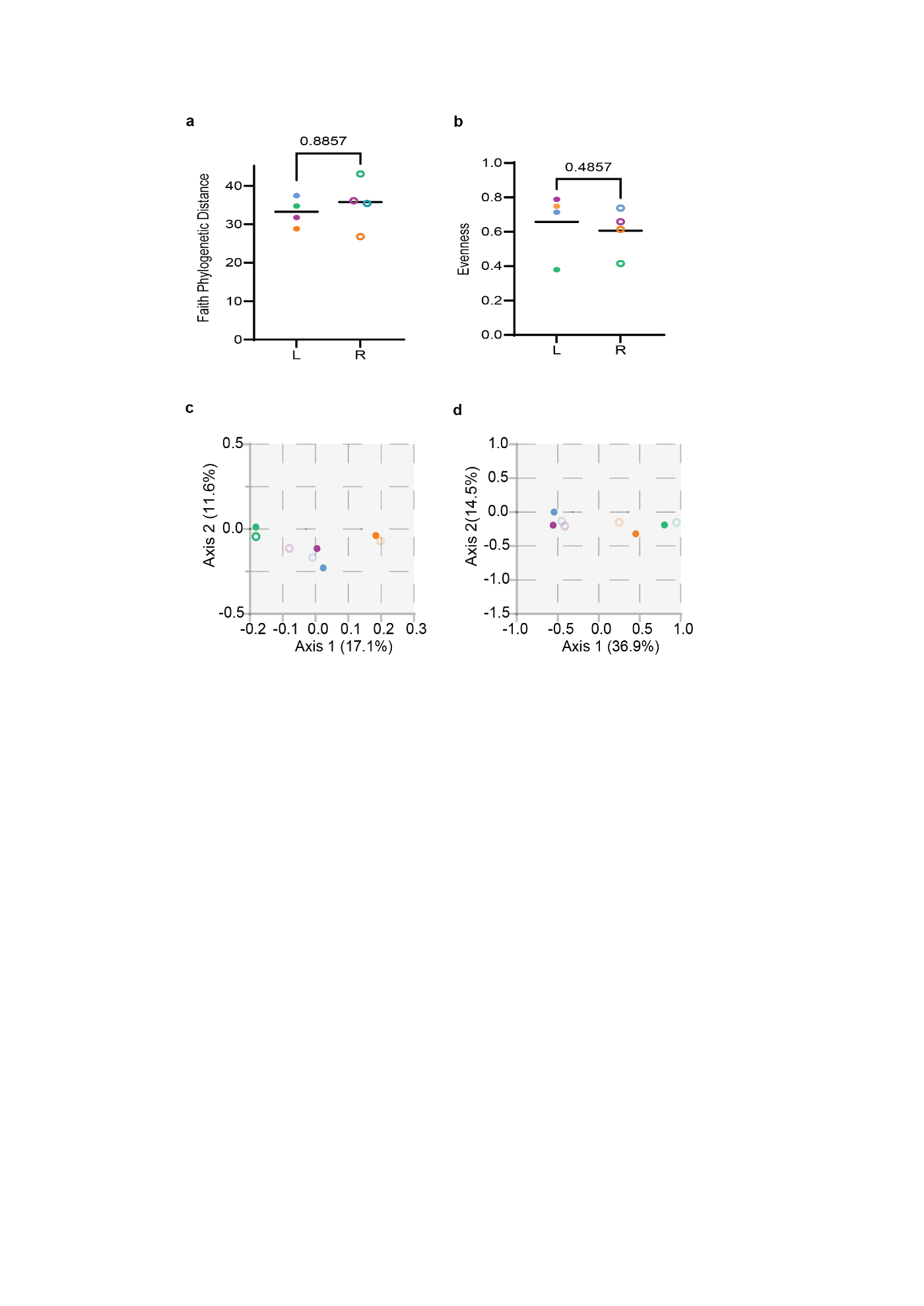

**Figure S1. Microbiome diversity is not different between matched Tonsil samples in DS enrollees. A**. Faiths Phylogenetic Diversity (FPD) **B.** Evenness **C.** PCOA analysis of Unweighted and Weighted UniFRAC **D.** Samples were compared by Single **(A-B)** Mann Whitney Test. P values are indicated on the graph. L: Left Tonsil, R: Right tonsil. Solid Circle: Left Tonsil. Open Circle: Right Tonsil. Colors are matched per donor.

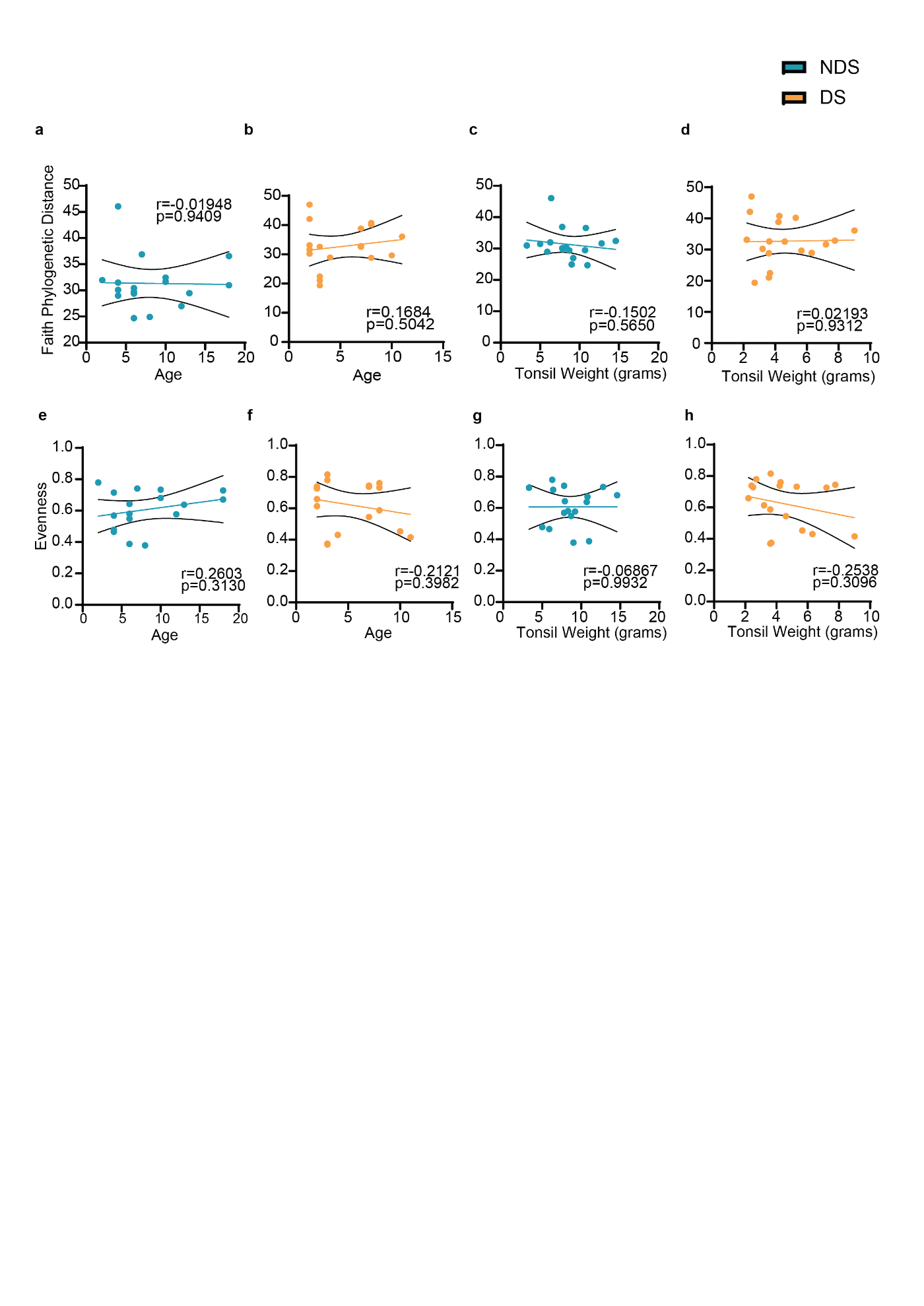

**Figure S2: No relationship between Tonsil microbiome Alpha Diversity with enrollee Age or Tonsil weight.** Spearmans correlation analysis of **A-D** Faiths Phylogenetic Distance or **E-H** Evenness or with enrollee Age (**A,B,G,H)** Tonsil Weight **(C-D)** or B.M.I. (**E-F)** in NDS (**A,C,E,G)** or DS (**B,D,F,H,)** Participants. DS: Orange. NDS: Teal. Circles: Each X,Y pair per enrollee. Lines: Linear regression. Dotted lines: 95% Confidence Interval.

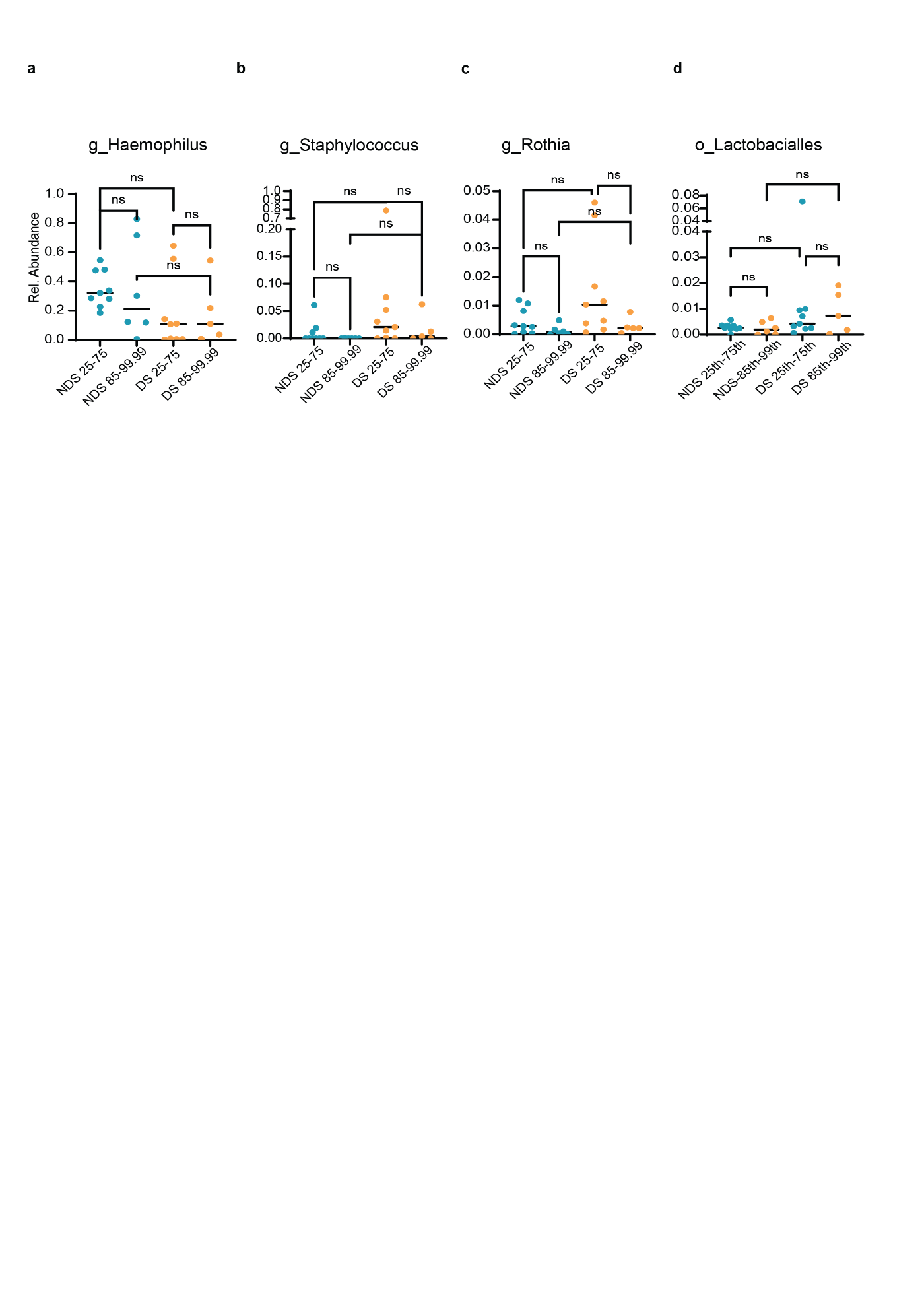

**Figure S3. No impact of BMI on differentially abundant organisms in DS. A-D** Abundance of **A.** Haemophilus, **B.** Staphylococcus, **C.** Lactobacialles and **D.** Rothia stratified by BMI per age percentile.DS (Orange) and NDS (Teal). Samples were compared by Single or **E-H** Kruskal-Walls , with Dunn correction for Multiple Comparisons*: p<0.05. ns: Nonsignificant. G_: genus. O_: Order
